# Conservation of co-evolving protein interfaces bridges prokaryote-eukaryote homologies in the twilight zone

**DOI:** 10.1101/067587

**Authors:** Juan Rodriguez-Rivas, Simone Marsili, David Juan, Alfonso Valencia

## Abstract

Protein-protein interactions are fundamental for the proper functioning of the cell. As a result, protein interaction surfaces are subject to strong evolutionary constraints. Recent developments have shown that residue co-evolution provides accurate predictions of heterodimeric protein interfaces from sequence information. So far these approaches have been limited to the analysis of families of prokaryotic complexes for which large multiple sequence alignments of homologous sequences can be compiled. We explore the hypothesis that co-evolution points to structurally conserved contacts at protein-protein interfaces, which can be reliably projected to homologous complexes with distantly related sequences. We introduce a novel domain-centred protocol to study the interplay between residue co-evolution and structural conservation of protein-protein interfaces. We show that sequence-based co-evolutionary analysis systematically identifies residue contacts at prokaryotic interfaces that are structurally conserved at the interface of their eukaryotic counterparts. In turn, this allows the prediction of conserved contacts at eukaryotic protein-protein interfaces with high confidence using solely mutational patterns extracted from prokaryotic genomes. Even in the context of high divergence in sequence, where standard homology modelling of protein complexes is unreliable, our approach provides sequence-based accurate information about specific details of protein interactions at the residue level. Selected examples of the application of prokaryotic co-evolutionary analysis to the prediction of eukaryotic interfaces further illustrates the potential of this novel approach.

**Significance statement:** Interacting proteins tend to co-evolve through interdependent changes at the interaction interface. This phenomenon leads to patterns of coordinated mutations that can be exploited to systematically predict contacts between interacting proteins in prokaryotes. We explore the hypothesis that co-evolving contacts at protein interfaces are preferentially conserved through long evolutionary periods. We demonstrate that co-evolving residues in prokaryotes identify inter-protein contacts that are particularly well conserved in the corresponding structure of their eukaryotic homologues. Therefore, these contacts have likely been important to maintain protein-protein interactions during evolution. We show that this property can be used to reliably predict interacting residues between eukaryotic proteins with homologues in prokaryotes even if they are very distantly related in sequence.

## Introduction

Cells function as a remarkably synchronized orchestra of finely tuned molecular interactions, and establishing this molecular network has become a major goal of Molecular Biology. Important methodological and technical advances have led to the identification of a large number of novel protein-protein interactions and to major contributions to our understanding of cells and organisms functioning (1–3). In contrast, and despite relevant advances in EM-microscopy (4–6) and crystallography (7, 8), the molecular details of a large number of interactions remain unknown.

When experimental structural data are absent or incomplete, template-based homology modelling of protein complexes represents the most reliable option (9, 10). Similarly to modelling of tertiary structure for single-chain proteins, homology modelling of protein-protein interactions follows a conservation-based approach, in which the quaternary structure of one or more experimentally solved complexes with enough sequence similarity to a target complex (the templates) is projected onto the target. Template-based techniques have provided models for a large number of protein complexes with structurally solved homologous complexes (11–14). Unfortunately, proteins involved in homologous protein dimers tend to systematically preserve their interaction mode only for sequence identities above 30-40% (15). For larger divergences, defining the so-called twilight zone (15, 16), it is not possible to discriminate between complexes having similar or different quaternary structures (15, 17, 18). As a consequence, the quality of the final models strongly depends on the degree of sequence divergence between the target and the available templates.

In contrast to more traditional approaches based on homology detection and sequence conservation, contact prediction supported by residues co-evolution (19–30) makes use of sequence variability as an alternative source of information (31). The analysis of residue co-evolution has been successfully applied to contact prediction at the interface of protein dimers (32–39), eventually leading to *de novo* prediction of protein complexes assisted by co-evolution (34, 35). In these methods, co-evolutionary signals are statistically inferred from the mutational patterns in multiple sequence alignments of interacting proteins. Co-evolution based methods have been shown to be highly reliable predictors of physical contacts in heterodimers, when applied to large protein families with hundreds of non-redundant pairs of interacting proteins (33–35, 40, 41). Unfortunately, these methods cannot be straightforwardly applied to the analysis of eukaryotic complexes where paralogs are abundant, making it very difficult to distinguish their interaction specificities. **In consequence, many eukaryotic complexes remain out of reach for both template-based homology modelling** (18) **and co-evolution guided reconstruction.** To address this eukaryotic “blind spot”, it is essential to identify biological constraints that have been conserved along very divergent evolutionary distances, that could be used for guiding the reliable projection of structural information from remote homologues.

We test the hypothesis that strong co-evolutionary signals identify highly conserved protein-protein contacts, making them particularly adequate for homology-based projections. From a structural modelling point of view, we test if and when co-evolution-based contact predictions can be projected to homologous complexes. In particular, we focus on the paradigmatic problem of contact prediction in eukaryotic complexes based on co-evolutionary signals detected in distant alignments of prokaryotic sequences. To this aim, we develop a novel, domain-centred protocol to detect co-evolving residues in multiple sequence alignments of prokaryotic complexes and evaluate their accuracy in predicting inter-protein contacts in eukaryotes. Our results show that **when the signal of co-evolution in prokaryotic alignments is strong, conserved inter-protein contacts in eukaryotes can be reliably predicted solely using prokaryotic sequence information.** These results provide the basis for the application of co-evolution to assist *de novo* structure prediction of eukaryotic complexes with homologues in prokaryotes even when they are highly divergent in sequence, a scenario where standard template-based homology modelling is unfeasible or unreliable.

## Results

### Benchmark dataset to study the interplay between co-evolution and structural conservation at protein interfaces

In order to investigate the relevance of co-evolution in the structural conservation at protein-protein interfaces among highly divergent homologues, we created a dataset of prokaryotic and eukaryotic domain-domain interfaces that integrates structural and co-evolutionary data at the residue level. We started from the complete dataset of heterodimeric Pfam (42) domain-domain interactions with known 3D structure (43). Co-evolutionary analysis of a protein interface requires a large set of paired sequences from the families of two interacting proteins in the complex (33–35). Distant evolutionary relationships can be often unveiled only at the level of domains (44): therefore, we devised a novel domain-centred protocol that enables the detection and the alignment of many conserved families of interacting domains (*Materials and Methods*, Fig. *S1A*). We searched for homologous sequences of the interacting domains in 15271 prokaryotic genomes and built a joint alignment by pairing domains in genomic proximity (34, 35). We used proximity in the genome to identify the existence of a specific physical interaction between two domains (45, 46). This protocol retrieved 559 cases of domain-domain pairs having 3D structural evidence of interaction in at least one prokaryotic or eukaryotic species and containing more than 500 sequences in the corresponding (non-redundant) joint alignment (Fig. 1B, *Materials and Methods*). For every domain-domain pair in this set, we computed co-evolutionary z-scores for all the inter-domain residue-residue pairs, that quantify the direct mutual influence between two residues (33) in different domains. Finally, we obtained the set of co-evolving inter-domain pairs of residues by retaining those pairs for which a strong co-evolutionary signal was detected (*Materials and Methods*). We classified each domain-domain interface as intra- or inter-protein (Fig. 1A) if the majority of paired sequences are codified within the same or different genes, respectively. 401 out of 559 domain pairs were classified as intra-protein and 158 as inter-protein (Fig. 1B and *S1B*).

**Fig. 1.**
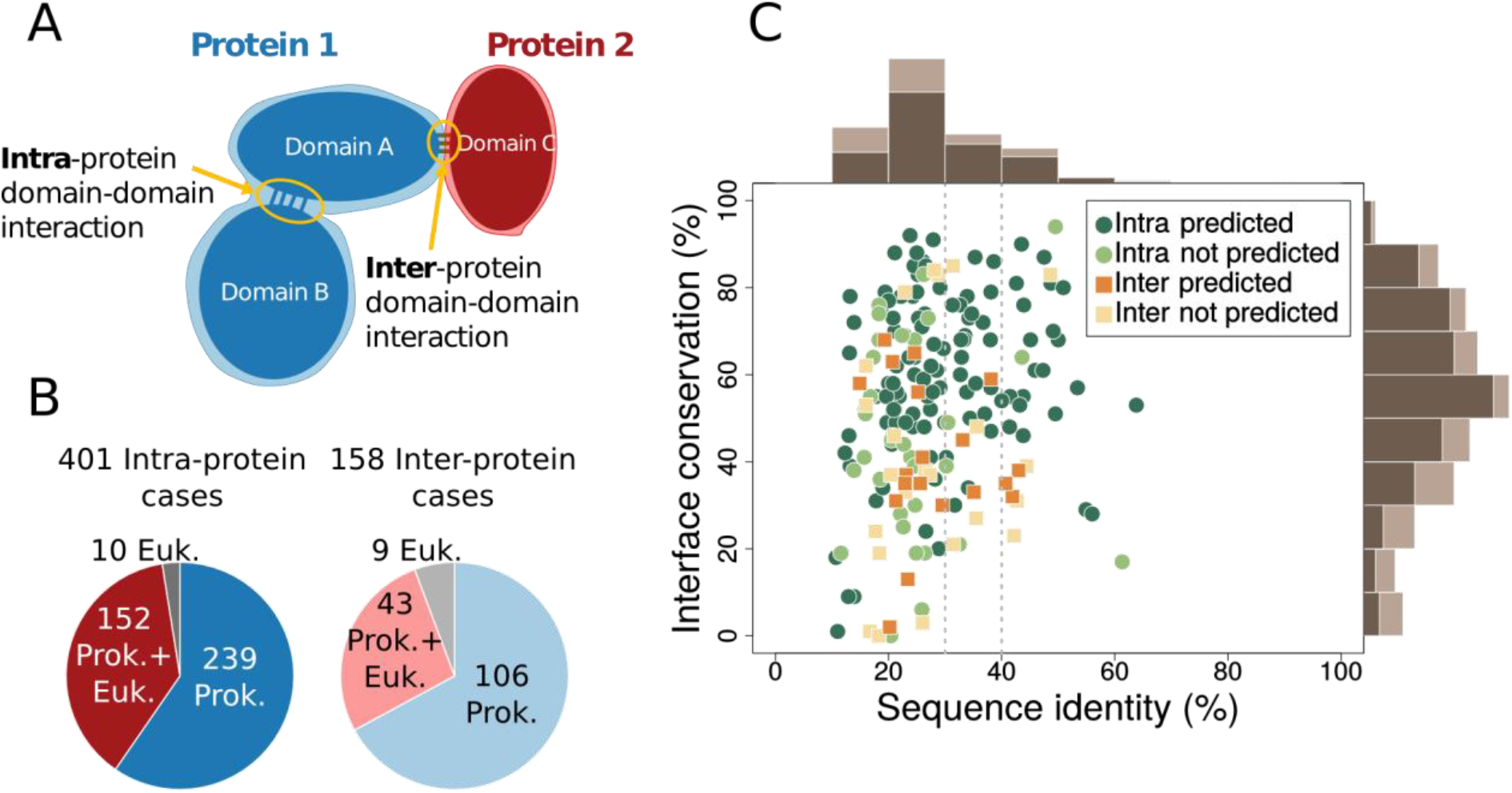
Summary of the co-evolutionary/structural dataset generated by our protocol. **(A)** Two kinds of domain-domain interactions discussed in the text: in intra-protein cases the two domains are codified within the same gene; in inter-protein cases they are found in different genes. **(B)** Dataset composition according to inter- or intra-protein classification and the availability of 3D structure in prokaryotes and eukaryotes. **(C)** Percentages of interface conservation and sequence identities for 152 intra-protein cases and 43 inter-protein cases. Interface conservation was calculated as the proportion of contacts in prokaryotic interfaces that are also in contact in eukaryotes. Sequence identities were calculated as the proportion of identical amino acids between the best aligned prokaryotic PDB sequence and the best aligned eukaryotic PDB sequence (SI Text). A domain-domain interaction was labelled as predicted when at least an inter-domain pair of residues was classified as co-evolving (Materials and Methods). Marginal histograms computed on the whole set of cases are in light brown; marginal histograms for predicted cases are in dark brown.

We first classified every 3D structure for each domain-domain interaction as prokaryotic or eukaryotic (*SI Text*). In order to deal with conformational variability in the available experimental structures, we used two different definitions for the set of contacts forming each domain-domain interface (see *Materials and Methods*): 1) we defined a *comprehensive interface* by merging all the inter-domain contacts (defined as residues closer than 8 Å, see *Materials and Methods*) extracted from all known homologous structures. This definition incorporates information from different biological (e.g. conformational changes) and methodological scenarios 2) we selected the complex that best aligned to the Pfam profile and defined the corresponding contacts as the *representative interface*. A comprehensive and a representative interface were computed separately for each domain-domain pair and for both eukaryotic and prokaryotic structures. When not specified otherwise, we will refer to the results obtained from the analysis of comprehensive interfaces; however, all the analyses were performed in parallel for the representative complexes with similar results. All the collected data were integrated in a dataset (Fig. 1) of 559 domain interactions with their inter-domain co-evolving residues and their corresponding prokaryotic and/or eukaryotic structural interfaces.

Our dataset includes 43 inter-protein and 152 intra-protein cases (Fig. 1B) with structure in both prokaryotes and eukaryotes. For these cases, we quantified the structural interface conservation as the proportion of prokaryotic contacts that are also in contact in their homologous sites in eukaryotes. In this subset, 66% (129 out of 195) of the cases corresponds to sequence identities below 30%. Complexes with sequence identities below 30-40% have highly variable values of interface conservation, and conserved interfaces cannot be identified using sequence identity alone (Fig. 1C, Fig. *S1C* for representative interfaces). This variability reflects the difficulties associated to accurate template-based homology modelling in the twilight zone. In our dataset, a naive extrapolation of contacts from prokaryotes to eukaryotes would result in highly unreliable predictions, due to the large divergences. This set of homologous interfaces provides the basis for investigating the structural conservation of co-evolving residues between prokaryotic and eukaryotic interfaces even at large sequence distances.

### Co-evolving residues identify structurally conserved contacts at protein interfaces

We detected strong co-evolutionary signals in 20 out of 43 inter-protein cases (and in 121 out of 152 intra-protein cases). The proportion of cases with predictions (strong co-evolutionary signals) is higher when the structural interface conservation is larger (Fig. 1C). This suggests that co-evolution is indicative of a greater structural conservation. To gain further insight, we studied the relationship between structural interface conservation and the degree of co-evolution detected in each case. To this aim, we calculated a score, called interface coupling, by averaging the z-score of the five strongest inter-domain co-evolving pairs (39). As shown in Fig. 2A, the level of interface coupling determines a lower bound for interface conservation, *i.e*. the stronger the interface coupling, the higher the minimal interface conservation observed in our dataset.

**Fig. 2.**
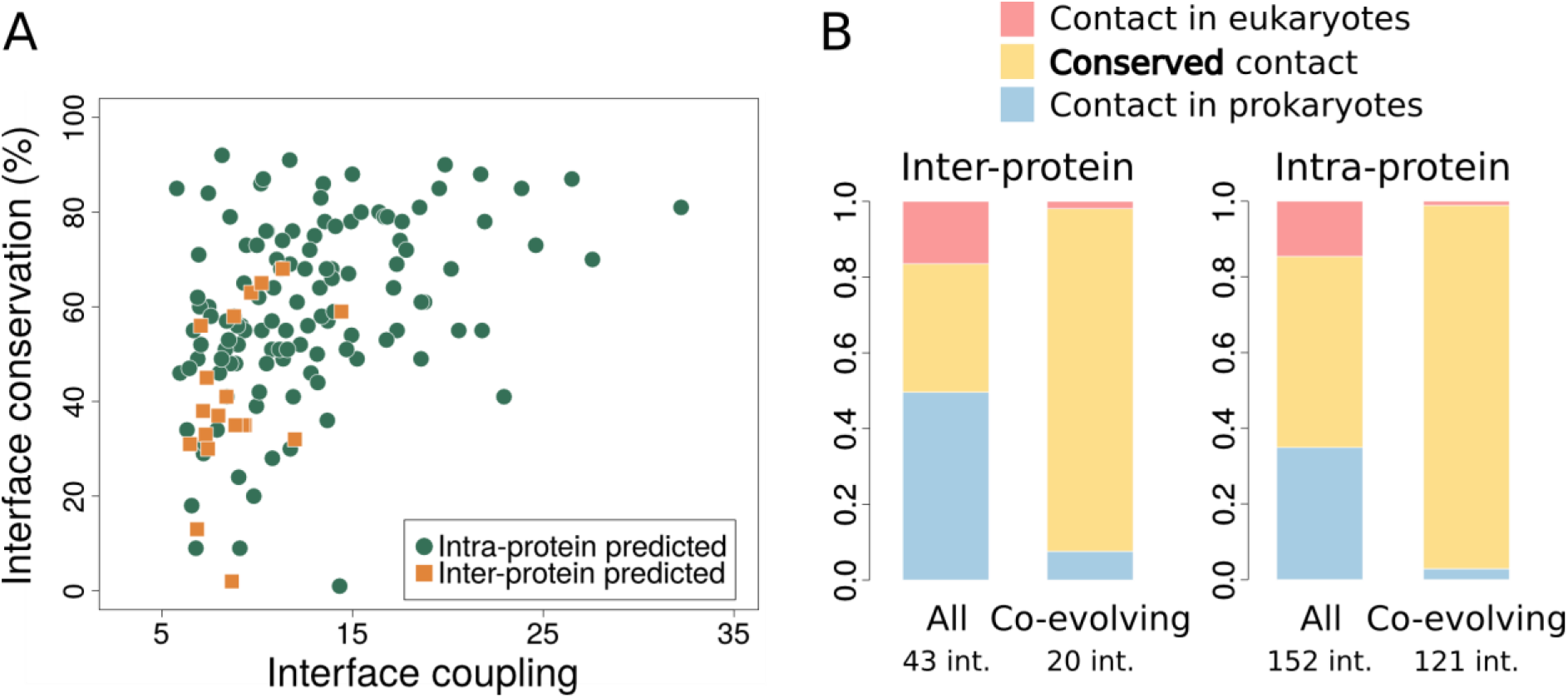
**(A)** Relation between interface structural conservation (defined as in Fig 1B) and interface coupling (the average z-score of the five strongest inter-domain co-evolving pairs) for 20 inter-protein and 121 intra-protein domain-domain interactions with contact predictions (i.e. strong co-evolutionary signals) and structurally solved prokaryotic and eukaryotic homologous complexes. **(B)** Proportion of conserved contacts at the homologous sites of prokaryotic/eukaryotic complexes, computed from the total set of contacts and the subset of co-evolving contacts, and for inter- and intra-protein cases. Blue: contacts found in a prokaryotic complex and not in the homologous eukaryotic complex. Red: contacts found in a eukaryotic complex and not present in the homologous prokaryotic complex. Yellow: contacts shared by prokaryotic and eukaryotic complexes. 48 out of 52 co-evolving contacts in inter-protein complexes and 1039 out of 1070 co-evolving contacts in intra-protein complexes are shared by prokaryotes and eukaryotes.

A comparison between homologous sites in eukaryotic and prokaryotic structures clearly reveals that **pairs of residues that are co-evolving and in contact in prokaryotes** (inter-protein: 52 contacts out of 56 co-evolving pairs; intra-protein: 1070 contacts out of 1107 co-evolving pairs) **are systematically found in contact in the 3D structures of the corresponding eukaryotic homologues** (Fig. 2B). This effect is highly significant when compared to the proportion of prokaryotic contacts shared with a eukaryotic homologue expected by chance (p-value < 10^−10^, one-tailed Fisher exact test for both inter-protein and intra-protein cases, *SI text*) and it is robust to different definitions of co-evolution and contacts (Fig. *S2A* *and* *B*). The analysis of representative interfaces leads to the same conclusion (Fig. *S2C* *and* *D*). Remarkably, focusing on the difficult cases in the twilight zone (less than 30% sequence identities, 29 inter-protein and 100 intra-protein) we also found a highly significant enrichment in conserved co-evolving contacts (Fig. *S3*, p-value < 10^−6^ one-tailed Fisher exact test for both inter-protein and intra-protein cases, *SI Text*).

In details, the proportion of inter-protein contacts conserved in prokaryotic and eukaryotic interfaces (34%) increases up to 92% (48 conserved contacts out of 52 co-evolving contacts) for pairs of co-evolving residues (Fig. 2B). Interestingly, 3 out of the 4 co-evolving pairs that apparently are not conserved correspond to residue pairs that are spatially close in the eukaryotic structure (less than 10 Å). For the cases in the twilight zone, 23 out of 24 co-evolving contacts are conserved in eukaryotes and the remaining pair is at 8.1 Å in eukaryotes. Intra-protein interfaces follow the same trend: the proportion of conserved contacts goes from 50% to 94% for co-evolving pairs (Fig. 2B; 1039 conserved contacts out of 1070 co-evolving contacts). Again, we found that co-evolving contacts are highly conserved even for interfaces in the twilight zone (583 conserved out of 607). These results clearly prove that co-evolving contacts have been preferentially conserved during the course of evolution, validating our hypothesis that co-evolution identifies structurally conserved contacts. Moreover, when applied to co-evolving pairs of residues at prokaryotic interfaces, this property should allow to predict interface contacts in eukaryotic proteins, in a wide range of evolutionary distances, including the twilight zone.

### Contact prediction at eukaryotic protein interfaces

We assessed the precision of prokaryotic co-evolving pairs in predicting contacts in prokaryotic and eukaryotic structures for cases with predictions in structurally solved regions, both in prokaryotic and eukaryotic interfaces (19 inter-protein, 120 intra-protein). **The vast majority of these cases have a high precision in the two superkingdoms** (Fig. *S4*). Only 1 out of 19 inter-protein cases in prokaryotes (6 out of 120 in intra-protein) was predicted with a precision lower than 0.6 (Fig. *S4*). For eukaryotes, these numbers are only slightly higher with 2 out of 19 inter-protein (11 out of 120 in intra-protein; Fig. *S4*). The few additional cases with low precision found for representative interfaces are evenly distributed in prokaryotes and eukaryotes suggesting that they are not related with the projection procedure (Fig. *S4*). Most false positives occur in cases within the twilight zone with low structural interface conservation (Fig. *S5*). This low structural conservation could result in poorly aligned eukaryotic sequences. We evaluated the impact of alignment quality on the projection of contact predictions from prokaryotes to eukaryotes, by computing the averaged expected alignment accuracy for residues at the eukaryotic homologous sites of the prokaryotic interface (*SI Text*). Indeed, most of the cases with low quality predictions in eukaryotes but not in prokaryotes correspond to low quality sequence alignments, both for comprehensive (Fig. *S6A* *and* *B*) and representative interfaces (Fig. *S6C* *and* *D*).

As discussed above, the high reliability of co-evolution as a predictor of contacts in prokaryotes, and the preferential conservation of co-evolving contacts allows to predict contacts in eukaryotes without any prior structural information. In order to further assess this point, we quantified the quality of eukaryotic contact prediction for all cases in which a eukaryotic structure was available to check the resulting predictions (51 inter-protein and 162 intra-protein; Fig 1B). We detected 62 co-evolving pairs in 22 inter-protein cases (~ 3 predictions per case) and 1140 pairs in 124 intra-protein cases (~ 9 per case). **We found that the precision in eukaryotes is very high both in inter-protein (precision = 0.81, Fig. 3A) and in intra-protein cases (precision = 0.95, Fig. 3B)** and it is only slightly lower than the precision obtained in prokaryotes (Fig. 3C and D; precision inter-protein = 0.86; precision intra-protein = 0.95). We repeated the analysis after removing cases with low alignment quality, using a filter based on the pairs of co-evolving residues (*SI Text*). In line with the discussion in the previous paragraph, the results suggest that an *a priori* filter can detect cases in which projected predictions have a lower precision (Fig. *S6E* *and* *F*, Table *S1*).

**Fig. 3.**
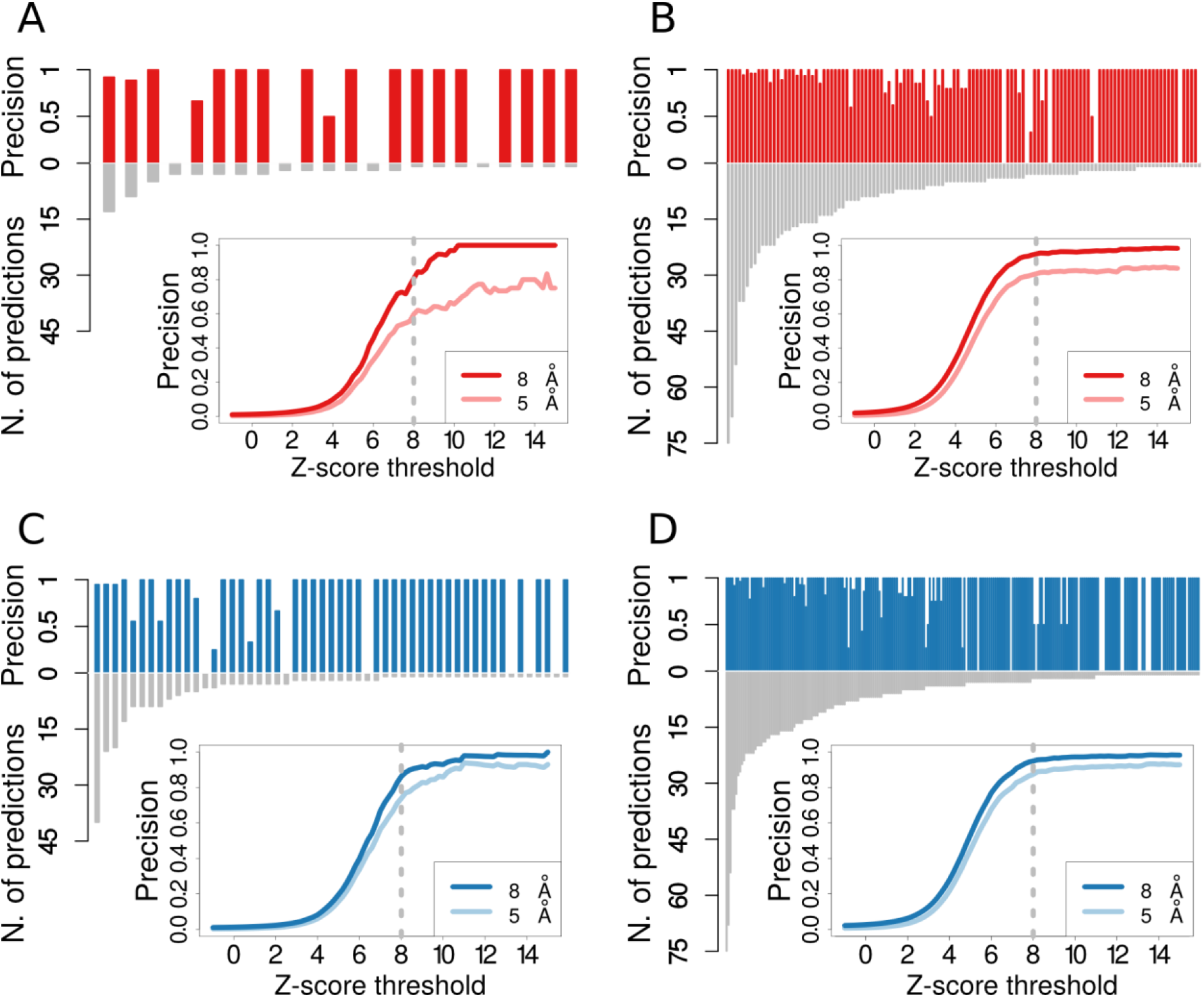
Precision and number of prediction for each of the 22 inter-protein **(A)** and 124 intra-protein **(B)** domain-domain interfaces in eukaryotes with at least one detected co-evolving pairs. Co-evolutionary z-scores were computed for all residue pairs for each domain-domain interface, and those pairs having a z-score larger than 8 were classified as co-evolving (Materials and Methods). Insets: the fraction of co-evolving pairs closer than 8 Å (dark blue line) and 5 Å (light blue line) as a function of the threshold for the z-score. The dashed grey lines highlight the reference z-score threshold (z-score = 8) for detection of co-evolution. We detected 62 co-evolving pairs for inter-protein interfaces (~ 3 prediction per case on average) with an average precision of 0.81 for a contact distance of 8 Å (and 0.6 at 5 Å), and 1140 pairs in the intra-protein case (~ 9 per case, on average) with a precision of 0.95 at 8 Å (and 0.83 at 5 Å). Precision and number of predictions for each of the 53 inter-protein **(C)** and 245 intra-protein **(D)** domain-domain interfaces in prokaryotes with at least one detected co-evolving pair. We obtained a precision of 0.86 at 8 Å and 0.74 at 5 Å for inter-protein domain-domain interfaces (respectively 0.95 and 0.87 in the case of intra-protein interfaces).

### Application to mammalian complexes

The pyruvate dehydrogenase complex, responsible of the catalysis of pyruvate to acetyl-CoA and CO_2_, is the complex in our dataset with the highest interface coupling in eukaryotes. Its E1 component forms a homo-dimer of hetero-dimers of its α and β subunits (47). The co-evolving contacts detected by our protocol are distributed over the interface between the two subunits and are well conserved in the eukaryotic interface. Among the 13 co-evolving pairs, 10 are in contact at the subunits interface of the branched-chain 2-oxo acid dehydrogenase remote homologue in the *Thermus thermophilus* structure (Fig. 4A) and are conserved in the human pyruvate dehydrogenase complex (Fig. 4B). Two out of three apparent false positives do actually correspond to contacts at the homo-dimer interface. These results show that co-evolution has been key in the conservation of quaternary structure in the pyruvate dehydrogenase E1 component.

**Fig. 4.**
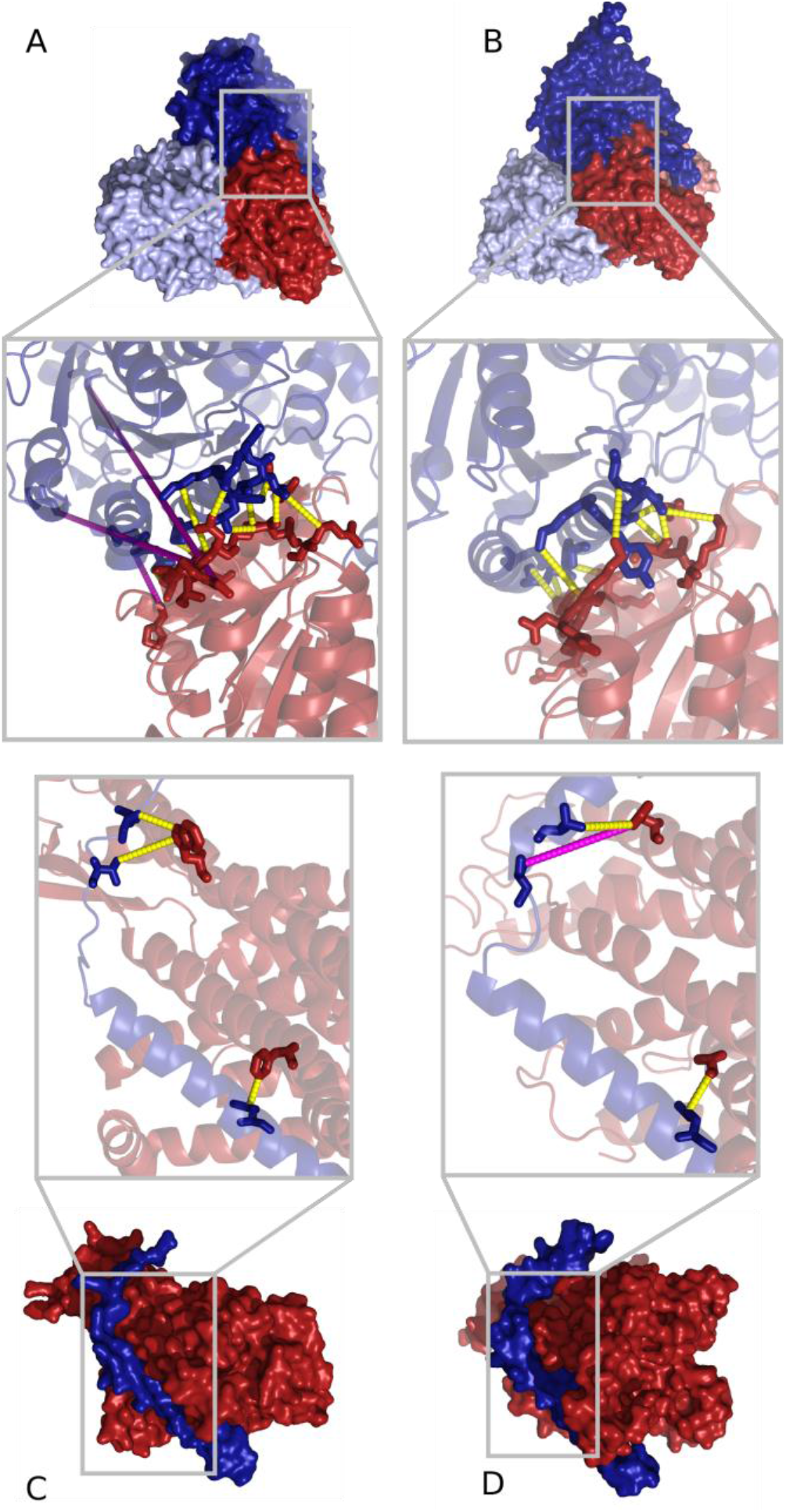
**(A and B)** E1 component of the branched-chain 2-oxo acid dehydrogenase in Thermus thermophilus (panel A, PDB code: 1UMB (70)) *and the human mitochondrial pyruvate dehydrogenase E1 component of the pyruvate dehydrogenase complex (panel **B**, PDB code: 3EXI (47)). Zoomed inset: co-evolving pairs of residues are shown as sticks and connected by dashed lines (yellow if they are in contact, magenta otherwise). 10 contacts out of 13 co-evolving pairs are detected at the Thermus thermophilus interface (panel A). 11 out of these 13 pairs can be mapped to the human complex where they are all in contact (panel B). **(C and D)** Protein transportation channels across bacterial plasma membrane SecYE in Thermus thermophilus X-ray structure of SecY (in dark red) in complex with SecE (in dark blue) (panel **C**, PDB code: 2ZJS* (71)). *Eukaryotic endoplasmic reticulum Sec61 in Canis lupus electron microscopy structure of Sec61 complex with its α subunit in dark red, γ subunit in dark blue (panel **D**, PDB code: 4CG7* (72)). The three co-evolving pairs (drawn as sticks) are in contact in Thermus thermophilus and two of them are conserved in the Canis lupus structure.

The translocon complex, one of the main mechanisms of transporting proteins across the membrane, is a good example of a conserved mode of interaction with very low sequence conservation. This family of complexes translocates proteins across the cytoplasmic membrane in bacteria (SecYEG complex) and across the endoplasmic reticulum in eukaryotes (Sec61 complex). The α and γ subunits of the mammalian Sec61 are homologous to the bacterial proteins SecY and SecE, respectively. Despite the low sequence identity between these proteins in *Thermus thermophilus* and *Canis lupus* (18.8% SecY/Sec61α and 10.5% SecE/Sec61γ), and a strong structural divergence of the domains, two out of three co-evolving contacts (Fig. 4C) have been conserved (Fig. 4D). In fact, seven out the nine residue pairs in the crystal structure for *Thermus thermophilus* with the highest co-evolutionary z-scores are in contact and six of them are structurally conserved in *Canis lupus*. The lower resolution (6.8 Å) of the available electron microscopy in *Canis lupus* introduces some uncertainty on the definition of the interface. Still, our predictions support the overall arrangement of the interaction given in this experimental structure and highlights the potential use of our approach to refine atomic details of cryo-EM experiments.

Among the 20 cases of inter-protein interfaces with structural information in both eukaryotes and prokaryotes and with strong co-evolutionary signals, we only detected one case where a strong co-evolutionary signal does not go together with a, at least partially, conserved interface: the interaction between the α and β subunits of the phenylalanyl tRNA synthetase (PheRS). PheRS catalyses the attachment of a phenylalanine amino acid to its cognate transfer RNA molecule. Despite important differences between the prokaryotic and the eukaryotic PheRS complexes (48), several homologous domains can be found in both the a subunit (core catalytic domain) and β subunit (B5 and B3/4 domains) between prokaryotes and eukaryotes (Fig. *S7A* *and* *B*). The co-evolutionary analysis of the interaction between the core catalytic domain and the B3/4 domain detects two co-evolving pairs located at the *Thermus thermophilus* interface (Fig. *S7C*). These pairs are no longer aligned in the human B3/4 domain due to an insertion in the *Thermus thermophilus* PheRS compared to the human cytosolic complex as proved from a structural alignment (Fig. *S7D*), and therefore cannot be projected to the corresponding interface. Notably, this interacting region in *Thermus thermophilus* is inserted just at one of the two turns where the human interaction takes place (Fig. *S7D*). Moreover, the α subunit also interacts with the B5 domain of the β subunit and the three co-evolving contacts at the prokaryotic interface are completely preserved in the human PheRS (Fig. *S7G* *and* *H*). This example illustrates that even after a drastic event, such as removal of a region at the interface in one of the interacting proteins, the remaining co-evolving residues can keep pointing to the real interfaces.

## Discussion

In this work we introduce and validate an important property of co-evolving contacts at protein interfaces: their propensity to be preferentially conserved at large evolutionary distances. This behaviour is confirmed by the analysis of co-evolving residues between domains in 15271 prokaryotic genomes and their homologous sites in 3D structures of eukaryotic complexes. This previously unrecognized aspect of the evolution of protein interfaces highlights the important role of co-evolving residues in maintaining quaternary structure and protein-protein interactions. As a first and important consequence of this property, we show that **contacts at eukaryotic interfaces can be predicted with high accuracy using solely prokaryotic sequence data**, both for protein-protein and for domain-domain interfaces. We tested these conclusions by analysing a large dataset of prokaryotic/eukaryotic interfaces with a novel, domain-centred protocol. We were able to predict contacts in inter-protein eukaryotic complexes with a mean precision > 0.8 (Fig 4, Table *S1*). This result is particularly relevant taking into account that this level of accuracy was attained for predictions of contacts in highly divergent complexes (sequence identities lower than 30%), where standard homology modelling is hardly useful. We have also shown that the few errors in these prokaryote-eukaryote projections are generally associated to cases with low structural conservation, that can be detected *a priori* by checking the alignment quality. Moreover, we extended this analysis to domain-domain contact predictions, showing that intra-protein interfaces exhibit even stronger co-evolutionary signals leading to an increased precision in contact prediction. The analysis protocol we propose relies on sequence data only. As a consequence, **our strategy can provide useful information on a protein interface both in remote homology-based complex reconstruction and when no structural template is available**, and it is inherently complementary to current methods based on the analysis of structural similarity (49, 50) or sequence similarity (10, 11, 14, 51) to a set of available templates.

The main obstacle to structural modelling of eukaryotic protein complexes by means of co-evolution based approaches is the need of a large number of homologous interactions to permit statistical analysis. Eukaryotic complexes present a paradoxical scenario: large families of eukaryotic proteins are the result of duplication-based expansions, but these duplications make uncertain which paralogues of one family interact with which ones of the other. In the future, improvements aimed to disentangle the network of paralogous interactions will be fundamental to deal with eukaryotic interactions (52–56). Our approach, based on preferential conservation, tackles this problem for proteins with prokaryotic homologues, by looking at very divergent, well populated and easy to couple pairs of interacting prokaryotic proteins. The resulting *projected* contact predictions represent a novel source of structural information that can be easily incorporated in integrative structural computational methods (57–61) or used to improve the scope of the successful methods that already incorporate co-evolutionary information from closer homologues (28, 34, 35, 62–64). At a more general level, these results indicate that co-evolving contacts have played a fundamental role in the evolution of interacting surfaces as structurally conserved anchor points.

## Materials and Methods

### Dataset and joint alignments

We extracted a list of 4556 hetero-dimeric pairs of interacting Pfam domains with solved 3D structures (3did database (43)). For each pair of Pfam domains, our protocol searched for proteins containing members of at least one of these two Pfam domain families in 15271 prokaryotic genomes using HMMER software(version 3.0) (65, 66). Two domains were paired if they were in the same protein, adjacent proteins or when both had no other paralogous in the corresponding genome. From this set of pairs, a joint alignment was built by aligning each domain to its corresponding Pfam profile. We next applied a stringent set of quality controls (*SI Text*) including alignment coverage (> 80%) and redundancy (< 80%). Insertions were removed by considering only residues that were assigned to match states of the HMM model. We retained 559 domain-domain interactions with a large number of non-redundant sequences (> 500) for further analyses. Each pair of Pfam domains was classified as intra- or inter-protein if the majority of paired sequences are codified within the same or different genes, respectively (Fig. *S1*).

### Calculation of co-evolutionary z-scores

We retrieved the co-evolutionary z-scores by performing a (multinomial) logistic regression of each position in the joint alignment on the remaining positions, a standard network inference strategy (67) that has already been adopted for the analysis of monomeric protein sequences (27) in combination with *l_2_* regularization. For each residue-residue pair we computed the Corrected Frobenius Norm score (27, 68), a measure of statistical coupling between residues, from the (symmetrized) estimates of the coupling parameters. Finally, these values were standardized to reduce heterogeneity between cases and used as co-evolutionary z-scores. An inter-domain pair of residues were considered as co-evolving when their co-evolutionary z-score was higher than a threshold value of 8 (*SI Text* for details).

### Interface definition and contact prediction evaluation

For a given pair of Pfam families, we retrieved, from the Protein Data Bank (PDB) (69), the biological unit for all the structures of complexes in which two members of the families are in physical contact. The PDB identifiers were retrieved from the 3did annotations. For structures with multiple biological units, we selected the one labelled as first. We extracted the PDB sequences and aligned them to their corresponding Pfam domains. We classified each PDB structure as eukaryotic or prokaryotic (*SI Text*). We defined a comprehensive and a representative interface in one or both superkingdom depending on the availability of at least one 3D structure in prokaryotes and/or eukaryotes. To that aim, for each pair of Pfam domains: 1) we recovered the inter-domain contacts in all PDB containing those Pfam domains. We used a distance of 8 Å between any heavy atom as the distance threshold for contacts (34, 35). Other contact definitions were used and are shown when appropriate 2) we mapped all PDB positions to their corresponding positions in the Pfam HMM profiles 3) we selected the most reliably aligned PDB (according to the alignment bitscores, see *SI Text*) as the representative complex in prokaryotes and eukaryotes 4) using the alignments of PDB sequences against both Pfam domains, we retrieved the set of PDBs with a 98% or higher percentage of sequence identity with respect to the representative complex. The representative interface is composed by the collection of contacts of the PDBs in this latter set, while the comprehensive interface is composed by all the contacts found in the PDBs containing the Pfam domains. Both interfaces were separately computed for eukaryotes and prokaryotes. Only pairs of inter-domain residues that were both aligned and having geometric coordinates in at least one PDB file were used to compute the precision of contact predictions. The precision of contact prediction was calculated as: Prec = TP/(TP+FP) considering co-evolving pairs of residues (z-score > 8) as positives.

## Acknowledgements

We thank F. Abascal and M.L. Tress for helpful discussions. Support for this work was provided by the Spanish Ministry of Economy and Competitiveness (MINECO) Project BFU2015-71241-R, co-funded by ERDF-EU. We are indebted to the NMR and crystallography researches, who have deposited structures in the PDB Databank, as well as to the DCA community for its insightful works.

## Supporting Information

### SI Text

#### Dataset

3did (version 06_2014 (43)) contains 8651 pairs of Pfam domains with solved 3D structures, 4556 of which are heterodimers. We run our protocol for these 4556 heterodimers Pfam pairs. For 559 pairs of Pfam domains, our protocol produced joint alignments with more than 500 paired domain sequences (see *Domain-domain pairing protocol section*). We classified each domain-domain interface as intra- or inter-protein if the majority of paired sequences are codified within the same or different genes, respectively. 401 out of 559 domain pairs were classified as intra-protein and 158 as inter-protein (Fig. 1B and *S1*). Using the corresponding PDB sequence we classified each structure as eukaryotic or prokaryotic (as explained in the next paragraph).

#### Prokaryotic/eukaryotic structure classification

Using annotations from 3did, we retrieved the PDB identifiers, chains and ranges of 37126 domain-domain interfaces with known structure in PDB. Using its taxonomy identifier (extracted from SIFTS (73) annotations), we classified each PDB chain as prokaryotic or eukaryotic by traversing up the NCBI taxonomy (74) tree, all the way up to the level of “Eukaryotes” or “Bacteria” or “Archaea”. Interfaces for which both interacting regions belong to the same chain (and are annotated in SIFTS) were classified correspondingly. Interfaces for which the two interacting regions belong to different PDB chains were classified when both chains were annotated in SIFTS in the same superkingdom (prokaryotes or eukaryotes, viruses are removed). Finally, we extracted their PDB sequences for the remaining unclassified structural interfaces and we searched them locally against a TrEMBL sequence database (75) using the blastp algorithm (version 2.2.18 (76)). Sequences having a similar sequence in the TrEMBL sequence database (percentage of sequence identity > 80%) were annotated following the TrEMBL phylogeny annotations, and the interface was classified when the two interacting regions were annotated and belonged to the same superkingdom (prokaryotes or eukaryotes, not viruses). The remaining 387 unclassified pairs were discarded.

#### Domain-domain pairing protocol

Given a pair of Pfam domains (version 27.0), for each domain in the pair, our protocol searched for proteins containing a member of the corresponding Pfam family through hmmsearch (HMMER 3.0 package, (hmmsearch --noali --domtblout --cut_ga) across all the coding regions of the 15271 prokaryotic genomes (ensembl bacteria release 23). Our protocol implements two strategies to find interacting domains. The first strategy is by gene fusion and gene neighbouring. The protocol checks whether 1) (gene fusion) each hit found for the first domain has a hit of the second domain in the same protein. Overlapping sequences are paired when the overlap is less than 10% of the smaller domain, up to a maximum of 20 residues, the overlapping residues are removed in one of the two domains) 2) (gene neighbouring) each hit found for the first domain has a hit of the second domain in another protein in the same contig but closer than 300 base pairs of nucleotides (using the annotation provided by ensembl bacteria). If the same Pfam domain appears more than once in a protein, these hits are discarded to avoid noise as it is not possible to ensure that all of them interact with its partner Pfam domain. The second strategy is by uniqueness across the genomes: the protocol checks if both Pfam domains appear just once in a genome. Once the pairs are made, the sequences of each Pfam domain from each Pfam domain are separately aligned against the corresponding HMM Pfam profiles using the hmmalign command from HMMER (hmmalign options --trim --allcol). The two output alignments are merged considering only matches states to the HMM profile. Several quality controls and filters are applied: 1) if the same sequence has more than one pairing, all the pairs involving the sequence are removed to avoid any ambiguity 2) any pair where one of the two sequences have more than 20% of gaps are discarded 3) redundant paired sequences with greater than 80% sequence identity (considering both sequences as a whole) are filtered out using the cd-hit program (version 2 (77)) in order to reduce the phylogenetic bias.

#### Calculation of co-evolutionary z-scores

Let A_i_ be a discrete random variable representing the amino acids at position i in a joint alignment, and taking values from an alphabet of 21 letters corresponding to the 20 natural amino acids plus the gap state. We performed multiclass logistic regression of each position A_i_ on the remaining positions **A**_\i_ (67), using a pairwise model of interaction between amino acids. In this model, the conditional probability of observing an amino acid a_i_ at position i given all the others in a sequence a is given by: P(A_i_ = a_i_ | **A**_\i_ = **a**_\i_) ∝ exp [h_i_(a_i_) + Σ_j_J_ij_(a_i_, a_j_)], where a_i_ and a_j_ are the amino acids at positions i and j, respectively. The coupling parameters **J**, that regulate the interactions between amino acids at different positions in the alignment, were inferred by maximization of a *l*_2_-regularized version of the (log) conditional likelihood LL(**h,J**): **h**^*^, **J**^*^ = argmax_**h**,**J**_ [ LL(**h,J**) − λΣ_a_h_i_(a)^2^ − λΣ_j,a,b_J_ij_(a,b)^2^], where LL(**h,J**) = M^−1^Σ_s_log[P(A_i_ = a^s^_i_ | **A**_\i_ = **a**^s^_\i_)], **a**^s^ is the s-th sequence in the joint alignment, M is the total number of sequences and λ = 0.01 (68). The optimization was performed using an in-house Fortran code calling the MINPACK-2 (78) dvmlm routine, which implements a limited memory variable metric minimizer. The resulting coupling values were symmetrized and a score was computed for each pair of residues following a protocol proposed by Ekeberg *et al*. (68) (Corrected Frobenius Norm score). To reduce heterogeneity between cases, these values were standardized and used as co-evolutionary z-scores: z-score(i,j) = (score(i,j) - median) / (1.4826 MAD), where median and MAD are the median and the mean absolute deviation of the scores of all the pairs. An inter-domain pair of residues were considered as co-evolving when the corresponding co-evolutionary z-score was higher than a threshold value of 8, that resulted in a good trade-off between number and precision of predictions.

#### Interface definition

For a given pair of Pfam families, we retrieved the biological unit for all the structures of complexes in which two members of the families are in physical contact from the PDB. The PDB identifiers were retrieved from the 3did annotations. For structures with multiple biological units, the first one was selected. When a biological unit was not available, we analyzed the asymmetric unit. For NMR structures only the first model was considered. We extracted the PDB sequences and aligned them to their corresponding Pfam domains. We included in the analysis domain-domain interactions from the 3did annotations only when both domains are in the biological unit. When the biological unit contains more than one domain from the interacting Pfam families, we retrieved the shortest distance between all the possible domain-domain combinations for each pair of residues. Disordered and unaligned residues were not included in the analysis.

#### Calculation of eukaryote-prokaryote sequence identities and structural conservations

We classified each PDB structure as a eukaryotic or a prokaryotic structure. We defined one representative interface for these superkingdoms when at least one 3D structure is available (see *Materials and Methods*). For those pairs of Pfams domains with structure in both prokaryotes and eukaryotes, we calculate the percentages of sequence identity between the eukaryotic and the prokaryotic representative complex. We calculated the sequence identities for the two Pfam domains using the alignments of the PDB sequences against the Pfam profiles obtained with hmmalign. Following (15), we select the minimum of these two percentages of sequence identity as reference. The rationale behind is that the domains with the lower sequence identity would tend to be a better estimator of the divergence in the interaction (15). For every pair of interacting Pfams with at least one eukaryotic and one prokaryotic structure, we defined its structural conservation as the percentage of contacts at the (comprehensive or representative) prokaryotic interface whose homologous positions are also in contact in the corresponding eukaryotic interface.

#### Alignment quality measurement

In order to estimate how the alignment quality was affecting to our inter-domain couplings at the interface, we calculated the average minimum expected alignment quality of the eukaryotic sequences. We used the PDB sequences of the eukaryotic representative complexes (*Material and methods*) and considered the homologous sites at eukaryotic sequence of the residues at the prokaryotic comprehensive interface. We recovered the expected alignment quality per residue obtained by HMMER and considered gaps as positions with 0 expected quality. We averaged the expected alignment quality per residue for each one of the two domains, and select the minimum of these two averages. In order to avoid the requirement of a structurally solved interface at prokaryotes, we calculated an equivalent score based on the co-evolving residues. In this case, we are targeting cases where a low quality alignment is associated to those residues detected as interdependent.

#### Statistical analysis of preferential structural conservation of co-evolving contacts

The preferential structural conservation of co-evolving contacts was evaluated by applying Fisher’s exact test. The contingency table is based on two conditions: structurally conserved contact/non-conserved contact, co-evolving contact/non-co-evolving contact. We compared the proportion of expected conserved contacts with the proportion of conserved contacts in the presence of strong co-evolutionary signals. Given a contact threshold, a contact in the prokaryotic structure comprehensive (or representative) interfaces is structurally conserved if the corresponding homologous sites are also in contact in the eukaryotic comprehensive (or representative) interface, otherwise they are considered not conserved. Co-evolving pairs of residues are defined as those with a co-evolutionary signal above a given co-evolutionary threshold, otherwise, they are considered as non-co-evolving. The results of the test were: p-value ~ 10^−12^ for inter-protein cases (48 conserved contacts out 52); p-value ~ 10^−188^ for intra-protein cases (1039 out of 1070). A similar conservation is observed when the ten first predictions (with annotated distances in both prokaryotes and eukaryotes) for each case were considered (inter-protein: 118 out of 135, p-value ~ 10^−24^; intra-protein: 989 out of 984, p-value ~ 10^−130^). When considering only cases with lower than 30% sequence identity: inter-protein p-value ~ 10^−7^ (23 out of 24); intra-protein p-value ~ 10^−107^(583 out of 607).

#### Data availability

All the data discussed in this work is available at the website: cointerfaces.bioinfo.cnio.es

---

**Fig. S1.**
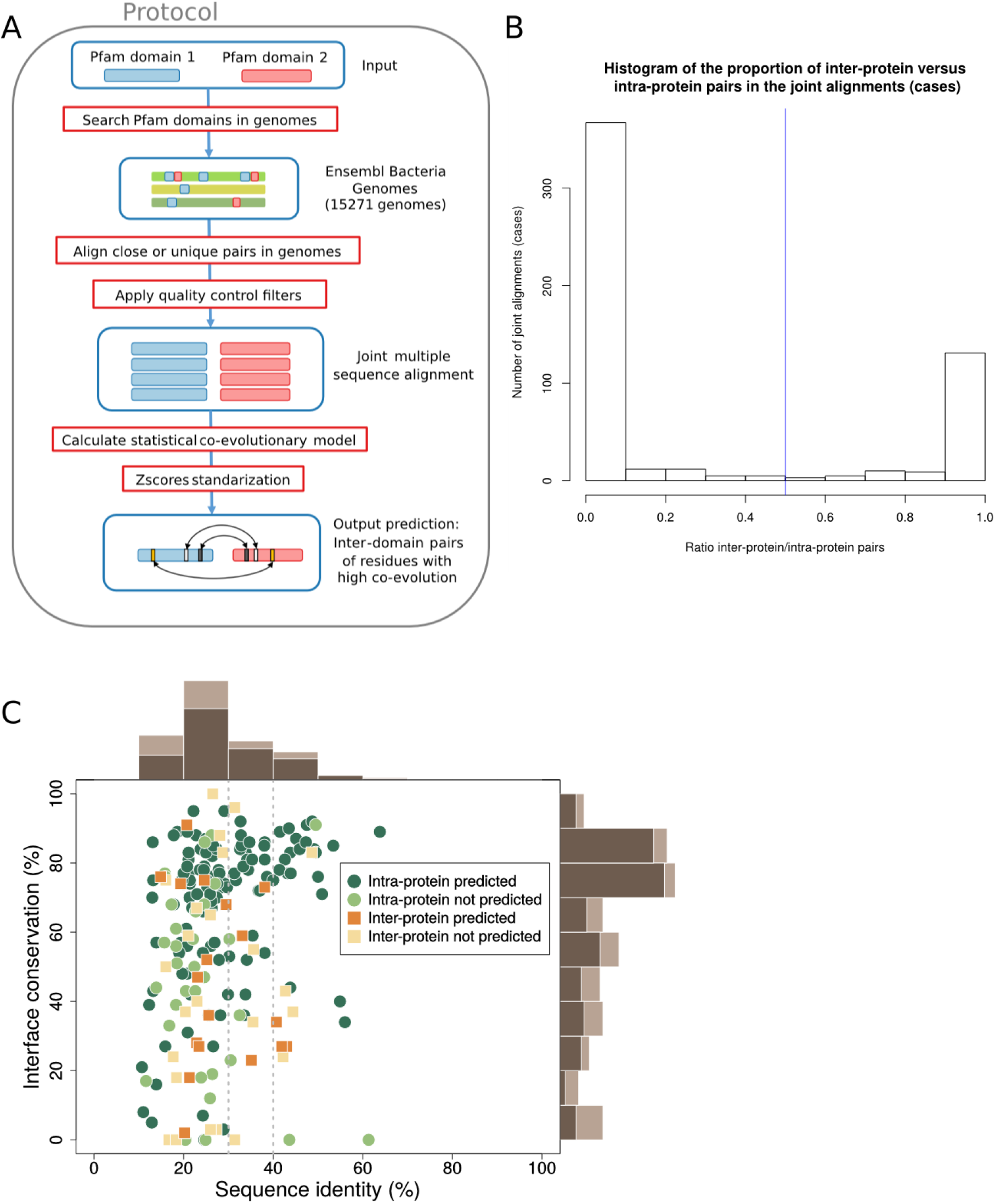
**(A)** Scheme of the co-evolutionary protocol developed. **(B)** Histogram of the ratio of inter-protein versus intra-protein pairs (0 means that all belong to the same proteins; 1 means that all domain pairs belong to different proteins) in the joint alignments. Cases of ambiguous classifications are rare: in *493* out of 559 cases, the paired sequences are predominantly (covering more than 90% of the pairs) found either in the same gene or in different genes. We classified a case as intra- or inter-protein when the ratio was lower or higher than 0.5, respectively. **(C)** Percentages of interface conservation and sequence identities for the 152 intra-protein and 43 inter-protein cases using representative interfaces. Interface conservation and sequence similarity were defined as described in Fig 1. Marginal histograms computed on the whole set of cases are in light brown; marginal histograms for predicted cases are in dark brown.

**Fig. S2.**
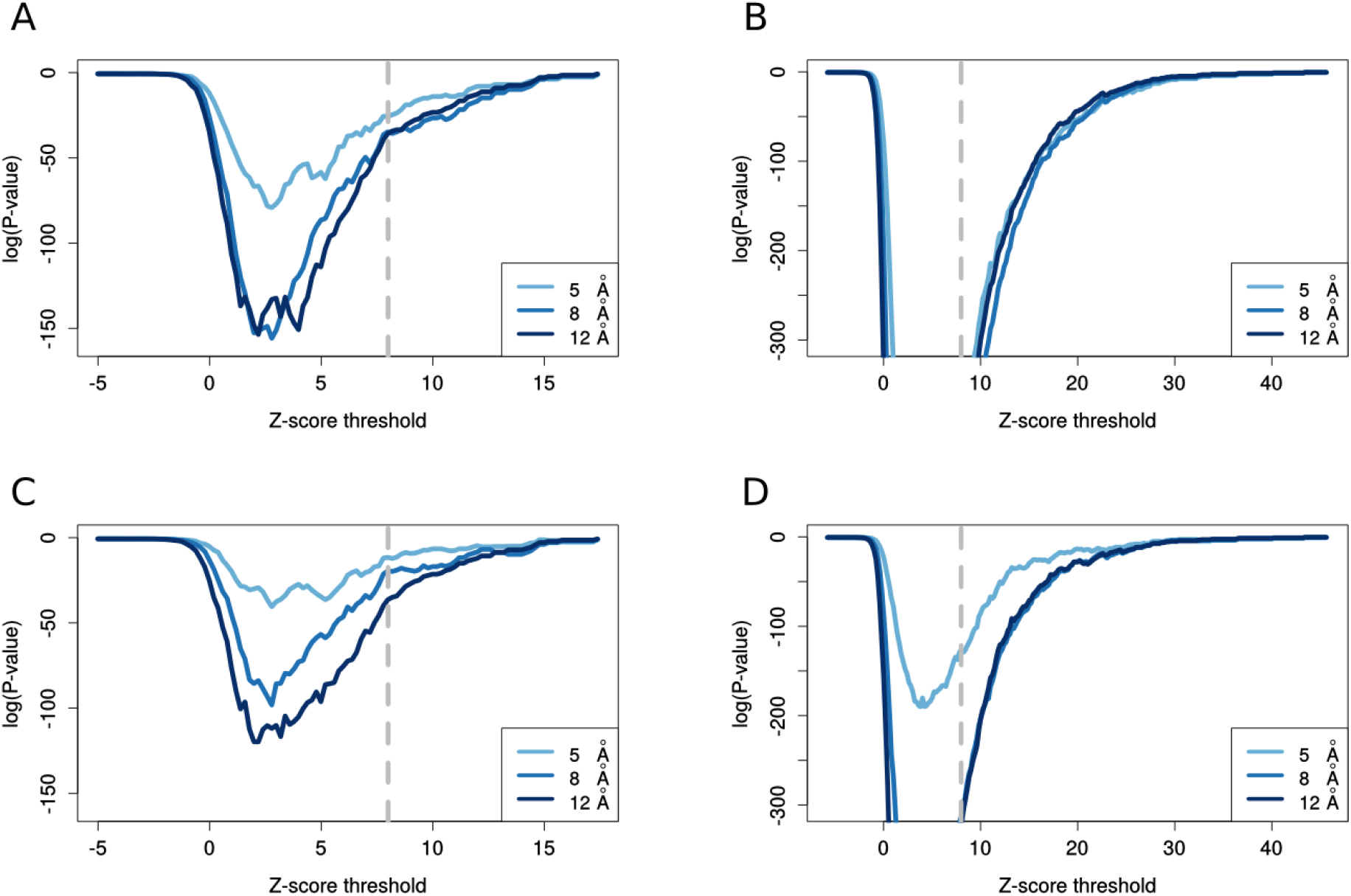
**(A and B)** One-tailed Fisher test p-value (*SI Text*) as a function of the z-score threshold (all co-evolving pairs with a z-score greater than the z-score threshold are considered) using comprehensive interfaces, for inter-protein **(A)** and intra-protein **(B)** cases. **(C and D)** One-tailed Fisher test p-value (*SI Text*) as a function of the z-score threshold using representative interfaces, for inter-protein **(A)** and intra-protein **(B)** cases. Note: P-values with log(p-value) < ~ −300 are beyond the floating point precision achievable using default R settings.

**Fig. S3.**
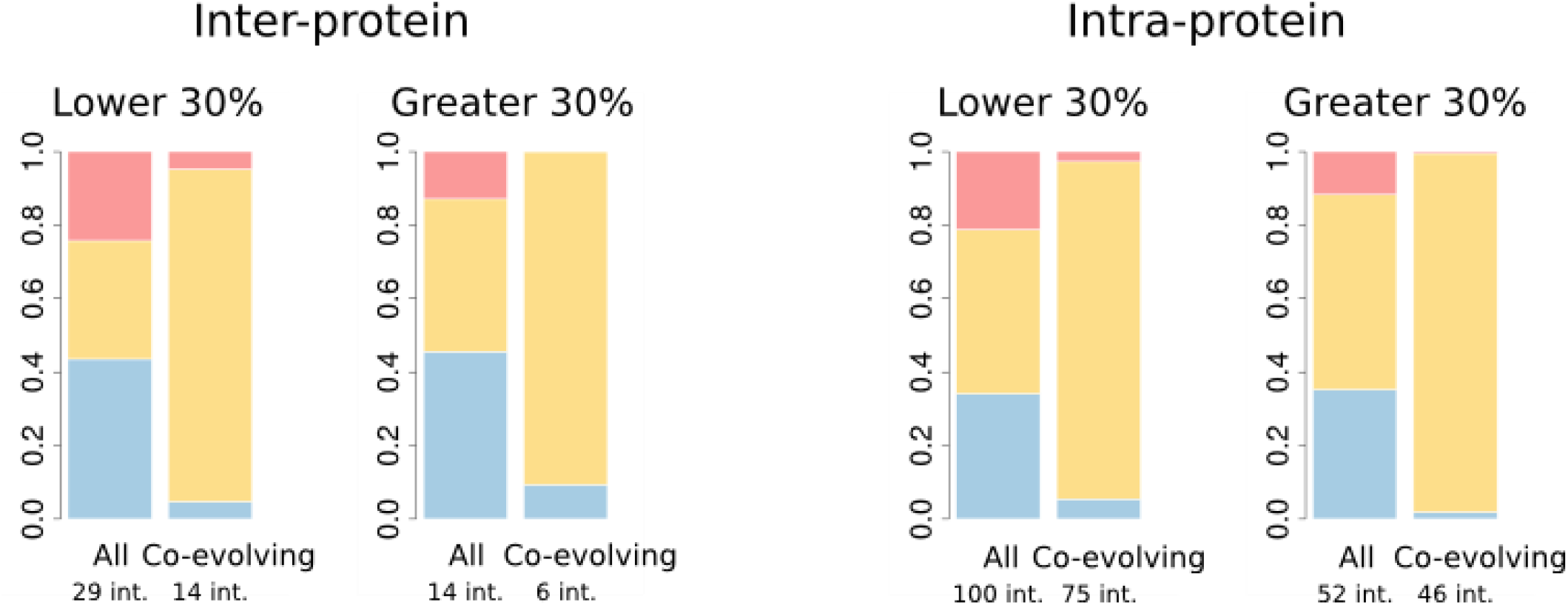
Proportion of contacts found only in prokaryotes, only in eukaryotes or conserved for all the contacts and for co-evolving contact. The cases are divided into two groups depending on whether the percentage of sequence identity (see *Materials and methods*) between prokaryotic and eukaryotic representative complexes are lower or greater than 30%.

**Fig. S4.**
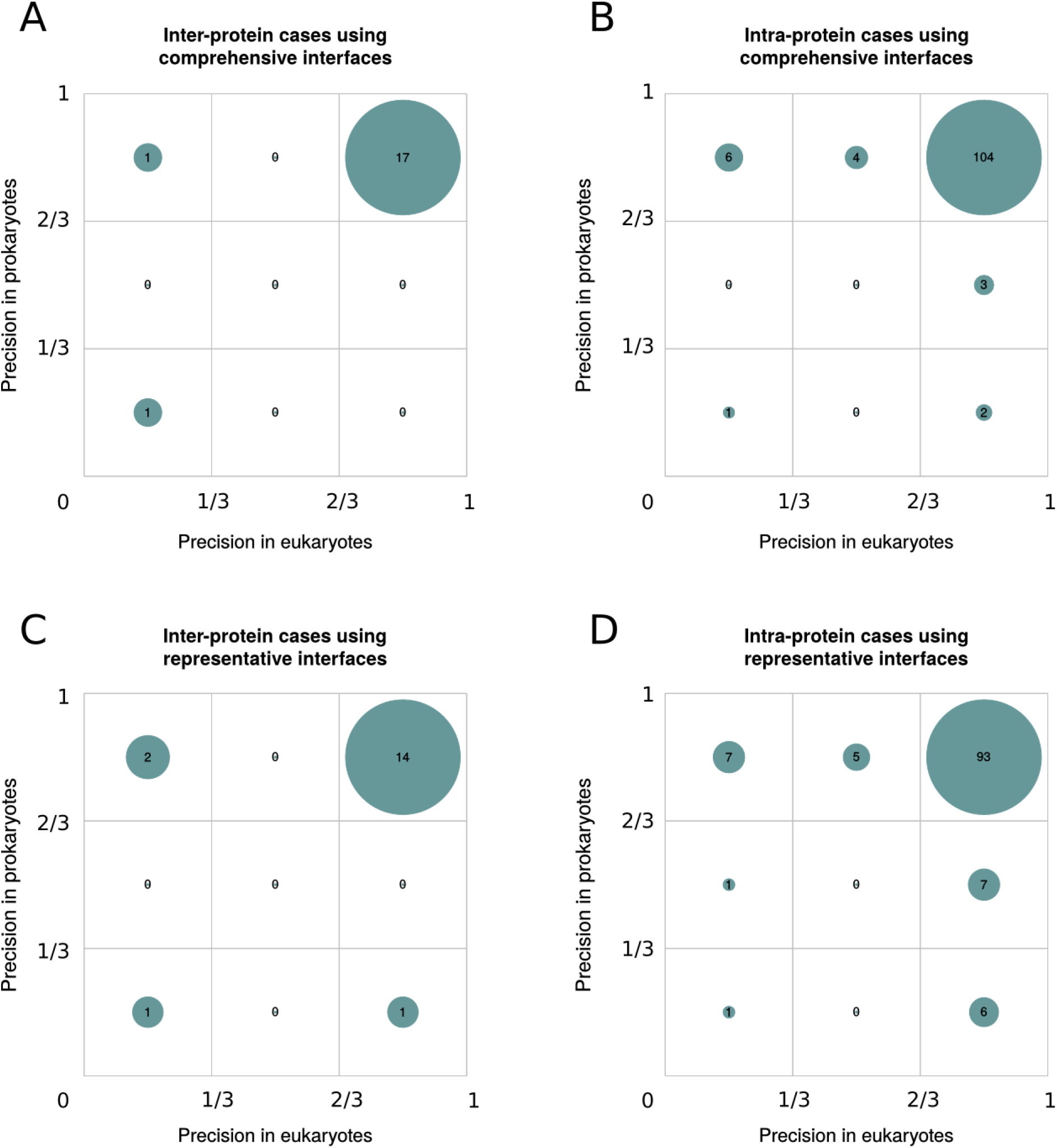
Precision comparison of contact predictions between prokaryotic and eukaryotic interfaces with 3D structures in both superkingdoms **(A)** Matrix of the number of inter-protein interfaces in three intervals of precision in both prokaryotes and eukaryotes using comprehensive interfaces. The areas of circles show the proportion of interfaces contained in the corresponding cell in comparison to the total number of interfaces in the matrix. **(B)** Matrix of prokaryotic and eukaryotic precisions of intra-protein interfaces using comprehensive interfaces. **(C)** Matrix of prokaryotic and eukaryotic precisions for inter-protein interfaces using representative interfaces. **(D)** Matrix of prokaryotic and eukaryotic precisions for intra-protein interfaces using representative interfaces.

**Fig. S5.**
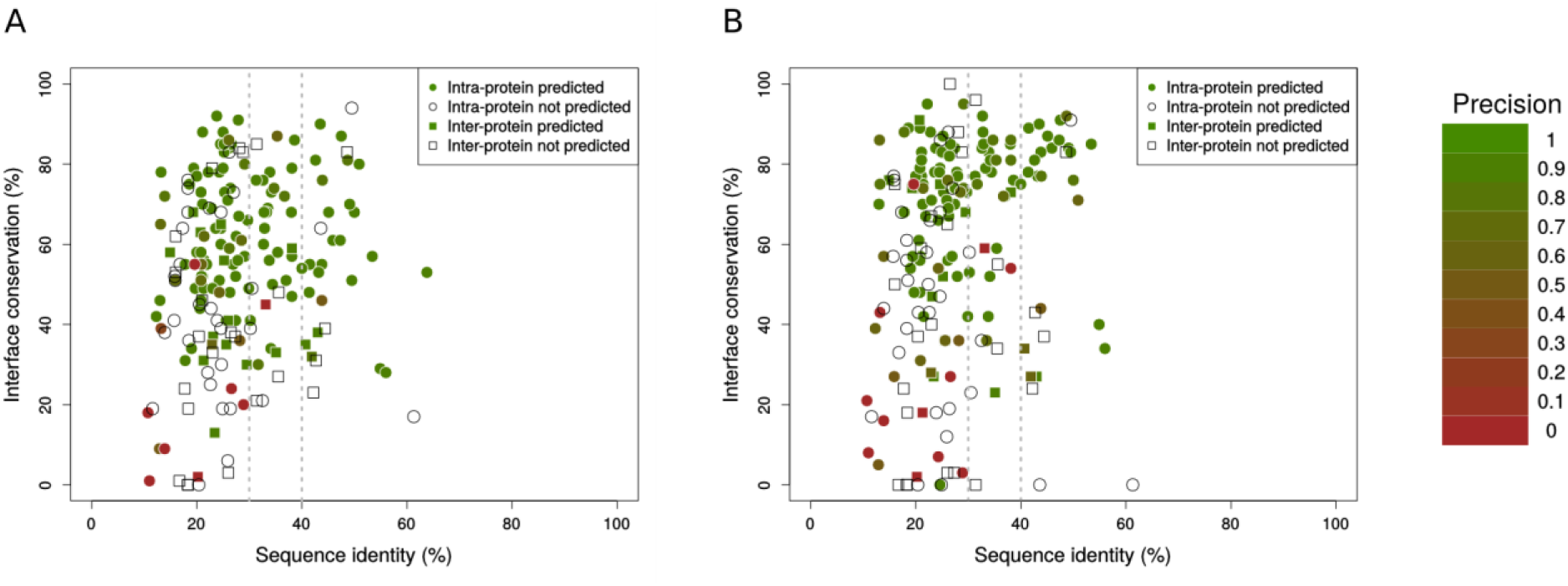
Each point represents a domain-domain interaction (inter-protein in squares, intra-protein in circles) with structure in both prokaryotes and eukaryotes, coloured by its contact prediction precision (white when there are not predictions). Percentages of sequence identity between the prokaryotic and eukaryotic representative complexes and interface conservation (see *SI Text*) are also shown. **(A)** Precision by case in eukaryotes as a function of the percentage of sequence identity and interface conservation using contacts from the comprehensive interfaces. **(B)** Precision by case in eukaryotes as a function the percentage of sequence identity and interface conservation using contacts from representative interfaces.

**Fig. S6.**
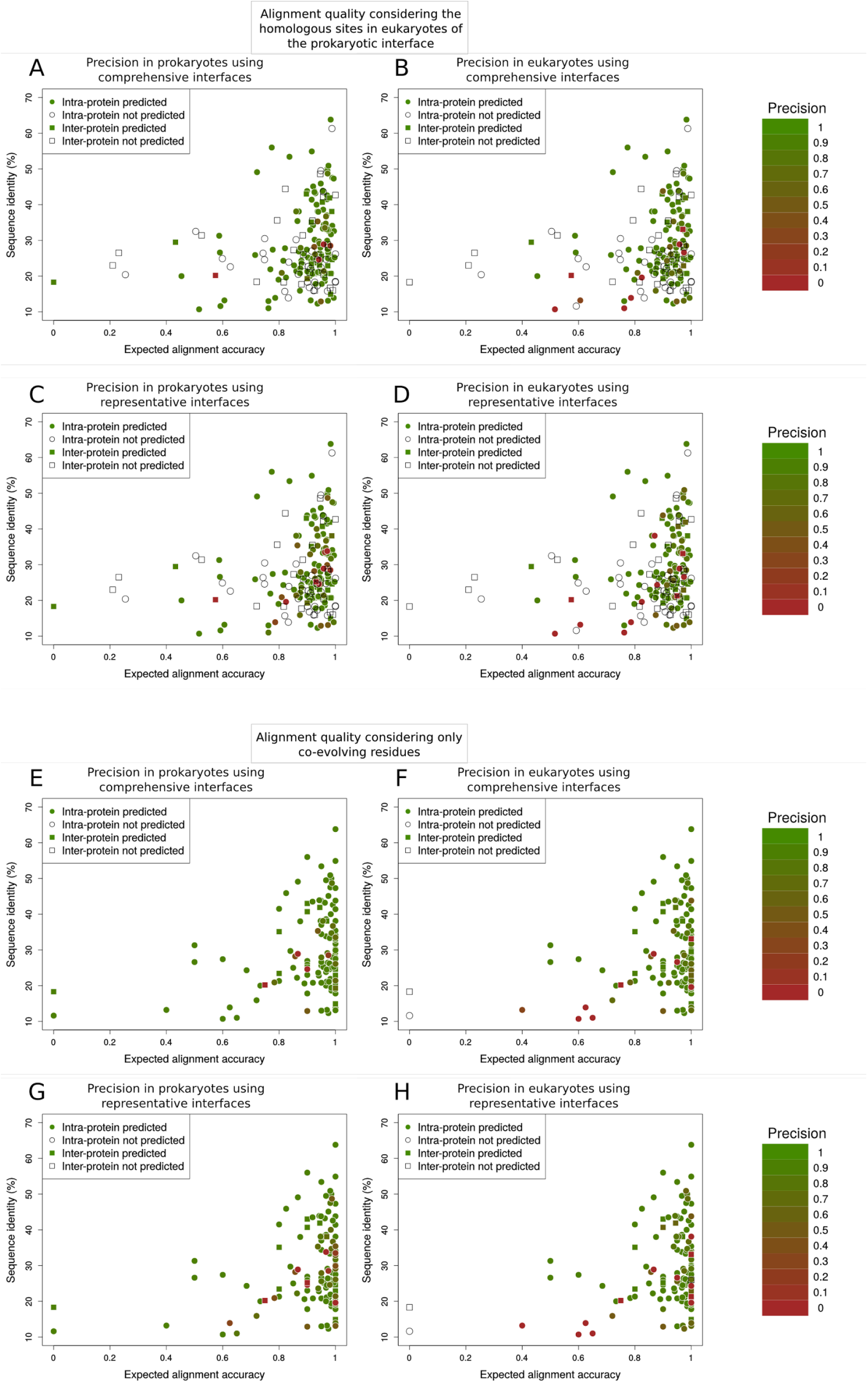
Quality alignment influence on the transference from prokaryotes to eukaryotes. Each point represents a domain-domain interaction (inter-protein in squares, intra-protein in circles) with structure in both prokaryotes and eukaryotes, coloured by its contact prediction precision (white when there are not predictions). Percentage of sequence identity between the prokaryotic and eukaryotic representative complexes and expected alignment accuracy (see *SI Text, Alignment quality measurement section*) are also shown. **(A-D)** The alignment quality has been computed considering the homologous sites in the eukaryotic sequences of the comprehensive prokaryotic interface, for comprehensive **(A-B)** and representative **(C-D)** interfaces. **(E-H)** The alignment quality has been computed considering only co-evolving residues (see *SI Text, Alignment quality measurement section*), for comprehensive **(E-F)** and representative interfaces **(G-H)**.

**Fig. S7.**
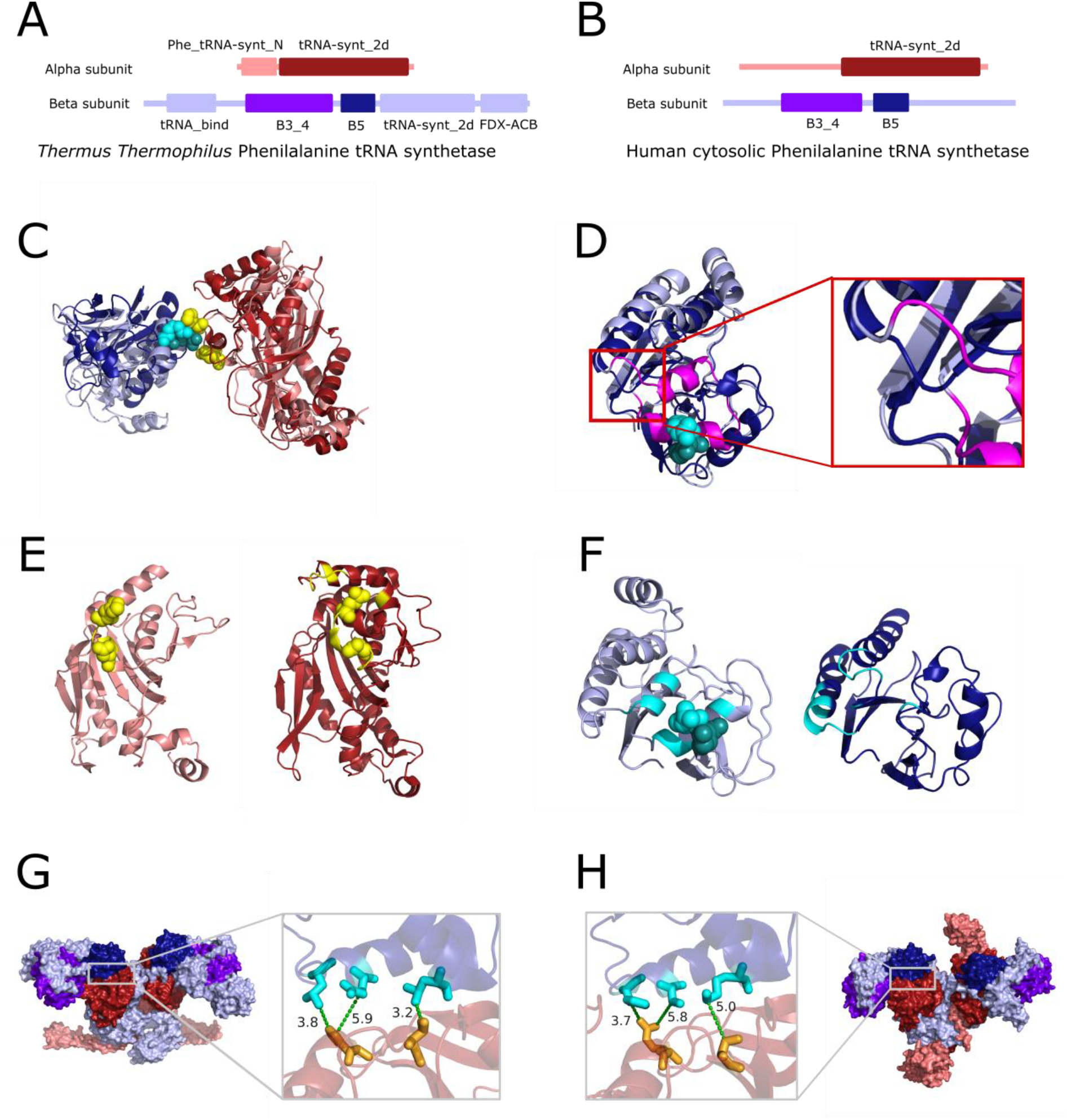
**(A and B)** Pfam domains found in *Thermus thermophilus* **(A)** and human cytosolic **(B)** phenylalanine tRNA synthetase are shown as rectangles. Pfam domains found in both eukaryotes and prokaryotes are drawn in darker colours, core catalytic domain (Pfam: tRNA-synt_2d) in α subunit in dark red, B5 and B3/4 (Pfam: B3_4) domains in β subunit in purple and dark blue respectively). **(C)** Superimposition of *Thermus thermophilus* (in light colors) PheRS structure and human PheRS (in dark colors) structure where the core catalytic domain (in reds) of both complexes have been structurally aligned. The B3/4 domains are shown in blues. Detected co-evolving pairs are shown as cyan spheres in the B3/4 domains and yellow in core catalytic domain. **(D)** Structural alignment of the *Thermus thermophilus* (in light blue) and human (in dark blue) B3/4 domain. The insertion in the *Thermus thermophilus* B3/4 domain is drawn in magenta. Zoom: The inserted region in detail. **(E)** Left: *Thermus thermophilus* core catalytic domain with interface residues in yellow and co-evolving residues in yellow spheres. Right: *Human* core catalytic domain with interface residues in yellow and projected co-evolving residues in yellow spheres. Despite the changes at the interface of the B3/4 domain, projected co-evolving residues at the human core catalytic domain are found at its interface. **(F)** Left: *Thermus thermophilus* B3/4 domain with interface residues in cyan and co-evolving residues in cyan spheres. Right: *Human* core catalytic domain with interface residues in cyan. **(G and H)** The three co-evolving pairs at the interface between the core catalytic domain and the B5 domain mapped to *Thermus thermophilus* **(G)** and human **(H)** structures are shown as sticks and connected by green dash lines together with the distance between the closest atoms in Å.

**Table S1.** Precision for contact predictions at eukaryotic interfaces. **(A)** Contact precision for inter- and intra-protein predictions and for both comprehensive and representative interfaces. For each dataset, the precision has been computed using the formula 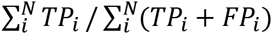, where TP_i_ and FP_i_ are respectively the number of true positive and false positives in the predicted contacts for the i-th interface. **(B)** Interface-averaged contact precision, computed as the average precision over the set of interfaces in each dataset: 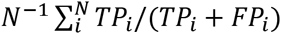. Standard errors were obtained by bootstrap resampling (10000 iterations with replacement).

| A. Precision |  |  |  |  |
| --- | --- | --- | --- | --- |
|  | Comprehensive |  | Representative |  |
|  | Before filter | After filter | Before filter | After filter |
| inter | 0.81 ± 0.05 | 0.91 ± 0.04 | 0.74 ± 0.06 | 0.83 ± 0.05 |
| intra | 0.95 ± 0.01 | 0.96 ± 0.01 | 0.90 ± 0.01 | 0.91 ± 0.01 |

**Table S1.** Precision for contact predictions at eukaryotic interfaces.
| B. Interface-averaged precision |  |  |  |  |
| --- | --- | --- | --- | --- |
|  | Comprehensive |  | Representative |  |
|  | Before filter | After filter | Before filter | After filter |
| inter | 0.77 ± 0.08 | 0.88 ± 0.06 | 0.69 ± 0.09 | 0.78 ± 0.08 |
| intra | 0.90 ± 0.02 | 0.93 ± 0.02 | 0.87 ± 0.02 | 0.89 ± 0.02 |

